# MICROSTRUCTURAL ORGANIZATION OF CORTICO-SUBCORTICAL FUNCTIONAL CONVERGENCE

**DOI:** 10.64898/2026.10.02.756182

**Authors:** Pragya Verma, Sakshi Singh, Shilpa Dang

## Abstract

The human subcortex integrates information from distributed cortical regions through functionally organized cortico-subcortical systems. Using the publicly available MICA-MICs multimodal MRI dataset of 50 healthy adults, we characterized the microstructural organization of functionally converging cortical regions. First, using resting-state fMRI., we identified cortical regions converging on subcortical targets, including thalamic, basal ganglia, and subcortical limbic structures. Then, using depth-resolved quantitative T1 profiles, we constructed a convergence-restricted microstructural profile covariance matrix, followed by diffusion-map embedding. The first microstructural gradient predominantly differentiated primary and association cortex, whereas the second gradient showed target-related differentiation beyond differences in primary versus association cortex. Cytoarchitectural analysis further revealed target-related differences in cortical composition. Together, these findings reveal complementary dimensions of cortico-subcortical functional organization, spanning broad cortical hierarchy, target-related microstructural differentiation, and cytoarchitectural composition.

## 1. INTRODUCTION

The human brain is organized across multiple spatial and functional scales, with distributed cortical regions interacting with subcortical structures to support diverse functions. Resting-state functional MRI (rs-fMRI) has revealed widespread cortico-subcortical functional interactions [1], [2]. Recent work has identified three major subcortical communities: thalamic nuclei (THA), basal ganglia regions (BGN), and subcortical limbic structures (SLN), that receive convergent inputs from distributed cortical regions [2].

Functional connectivity, however, does not directly characterize the microstructural properties of the cortical regions participating in this converging organization. Quantitative T1 (qT1) provides an in-vivo measure sensitive to regional microstructural variation and cyto/myeloarchitectonic organization [3]. Depth-dependent qT1 profiles can be compared across cortical regions using microstructural profile covariance (MPC), providing a measure of their microstructural similarity [4], [5]. Low-dimensional gradient approaches can further capture broad axes of cortical hierarchy, including the established transition between sensorimotor and transmodal/association cortex [6]. Microstructural gradients have also been linked to cortico-subcortical organization; for example, quantitative MRI gradients within the striatum have been related to cortical hierarchy and cortico-striatal microstructural covariation [7]. Further, cytoarchitectural classifications provide a complementary description of cortical microstructure based on differences in laminar differentiation and granularity [5], [8].

Despite above evidence, it remains unknown whether cortical regions converging on distinct subcortical communities exhibit systematic differences in their microstructural organization, and whether these differences extend beyond the established primary-to-association axis of cortical organization. Here, using the publicly available MICA-MICs multimodal MRI dataset [4], we integrated rs-fMRI-derived functional convergence with depth-resolved qT1 profiling to characterize the microstructural organization of cortical regions converging on THA, BGN, and SLN. We identified functionally converging cortical regions, constructed an MPC matrix restricted to these regions, and applied diffusion-map embedding to derive low-dimensional microstructural gradients. We then examined whether these gradients varied according to subcortical target and cortical type, accounting for the established primary-to-association axis of cortical organization [6], [7]. We additionally characterized cytoarchitectural composition to examine target-specific cytoarchitectural differences in cortical microstructure. We hypothesized that functionally converging cortical regions would exhibit systematic microstructural organization reflecting both cortical hierarchy and subcortical target-related differences.

## 2. MATERIALS AND METHODS

### 2.1. Participants and multimodal MRI data

We analysed multimodal MRI data from the MICA-MICs release [4], comprising 50 healthy adults (23 females; 29.54 ± 5.62 years) with resting-state fMRI and quantitative T1 imaging. For each subject, we employed the pre-processing pipeline on rs-fMRI data as indicated in the dataset study [4], including removal of the first five volumes, motion and distortion correction, ICA-FIX denoising, and motion-spike regression. Mean rs-fMRI time series were extracted from 54 subcortical parcels defined by the Tian atlas (27 per hemisphere) [9] and 400 cortical parcels from the Schaefer 7-network atlas (200 per hemisphere) [10]. For qT1, we used cortical profiles sampled at 14 equivolumetric depths and the corresponding subject-level microstructural profile covariance (MPC) matrices provided by the dataset.

### 2.2. Functional convergence analysis

For each subject, resting-state functional connectivity (FC) between the 54 subcortical and 400 cortical regions was estimated using multiple linear regression, with each region modelled against all remaining regions, as performed in [2]. Subject-level regression coefficients were entered into a second-level model to obtain group-level functional connections. Based on previously identified subcortical communities [2], we considered three target groups: thalamus (thalamic nuclei), basal ganglia (putamen, caudate, nucleus accumbens, and globus pallidus), and subcortical limbic structures (hippocampal and amygdala parcels). Group-level connections were thresholded at Bonferroni-corrected *p* < 0.05, and the resulting FC matrix was binarized. Cortical regions significantly connecting with at least one target group were defined as the functionally converging cortical set and were additionally classified as primary (PRI), including visual and somatomotor networks or as association (ASS) cortex, including dorsal and ventral attention, limbic, control, and default mode networks [10].

### 2.3. Microstructural profile covariance and gradient analysis

For each subject, the 14-depth qT1 profiles characterized intracortical microstructural variation across the 400 cortical regions [4]. Pairwise similarity between regional profiles was quantified using MPC. Subject-level MPC matrices were averaged to obtain a group-level MPC matrix, which was then restricted to the functionally converging cortical regions.

The convergence-restricted group-level MPC matrix was subjected to diffusion-map embedding using BrainSpace toolbox [11] and a normalized-angle kernel. Gradient components were ranked according to their normalized eigenvalue contributions; G1 and G2 were retained as the dominant gradients, while G3 and G4 were additionally examined for completeness. Each gradient provided a continuous coordinate value representing the position of each convergence-related cortical region along a microstructural axis.

G1 and G2 values were analysed separately using two-way ANOVA with *Target* (THA, BGN, SLN), *CortexType* (PRI, ASS) as factors. Significant effects involving multi-level factors were followed by Bonferroni-corrected pairwise comparisons (*p* < 0.05). The same model was applied to G3 and G4 as supplementary analyses. Cortical types were included to account for the established primary-to-association axis of global cortical microstructural organization, thereby distinguishing target-related differences from those attributable to broad cortical hierarchy [6].

### 2.4. Cytoarchitectural composition of cortical convergence regions

To characterize what aspects of cortical microstructural organization i.e., to distinguish regions showing convergence with different subcortical targets, we additionally examined their underlying cytoarchitectural composition, focusing on differences in laminar organization across cortical regions. The 400 cortical parcels of the Schaefer 7-network atlas were mapped to a published von Economo & Koskinas cytoarchitectural classification [8]. The classification comprises six cytoarchitectural types: koniocortical (Kon), eulaminate (Eul-III, Eul-II, Eul-I), dysgranular (Dys), and agranular (Agr), which capture differences in cortical laminar differentiation and granularity. For each of the cortical regions showing functional convergence, the proportion of vertices assigned to each cytoarchitectural type was calculated, yielding a six-element percentage profile for each cortical region.

The resulting cytoarchitectural profiles were analysed using a three-factor mixed repeated-measures ANOVA, with *CortexType* (PRI, ASS) and *Target* (THA, BGN, SLN) as between-region factors and *CytoType* (Kon, Eul-III, Eul-II, Eul-I, Dys, Agr) as a within-region repeated factor. The percentage assigned to each cytoarchitectural type was the dependent variable. Significant interactions were followed by Bonferroni-corrected pairwise comparisons (*p* < 0.05) to identify the specific cytoarchitectural types contributing to target- and cortex-type-related differences.

## 3. RESULTS

### 3.1. Functional convergence and microstructural covariance

Resting-state FC identified multiple cortical regions showing significant functional associations (p < 0.05, Bonferroni-corrected) with at least one of the three predefined subcortical target communities: THA, BGN, and SLN (Binary FC matrix shown in **Fig. 1A**). **Fig. 1D** illustrates the extent of cortical convergence across individual subcortical regions, with the number of converging cortical regions ranging from 0 to 6, represented by node size. Across the converging cortical regions, 60% showed convergence with SLN, 30% with BGN, and 10% with THA, indicating a greater concentration of cortical convergence toward the subcortical limbic community. The convergence-related cortical regions comprised both primary and association cortical areas. The target-specific convergence sets differed in their cortical composition, with THA convergence predominantly involving primary cortical regions (78%), whereas BGN and SLN convergence involving association cortex (68% and 82%, respectively). The dominant cortical networks also differed across target communities, with VIS, VAN, and DMN predominating for THA, BGN, and SLN convergence, respectively.

**Fig. 1.**
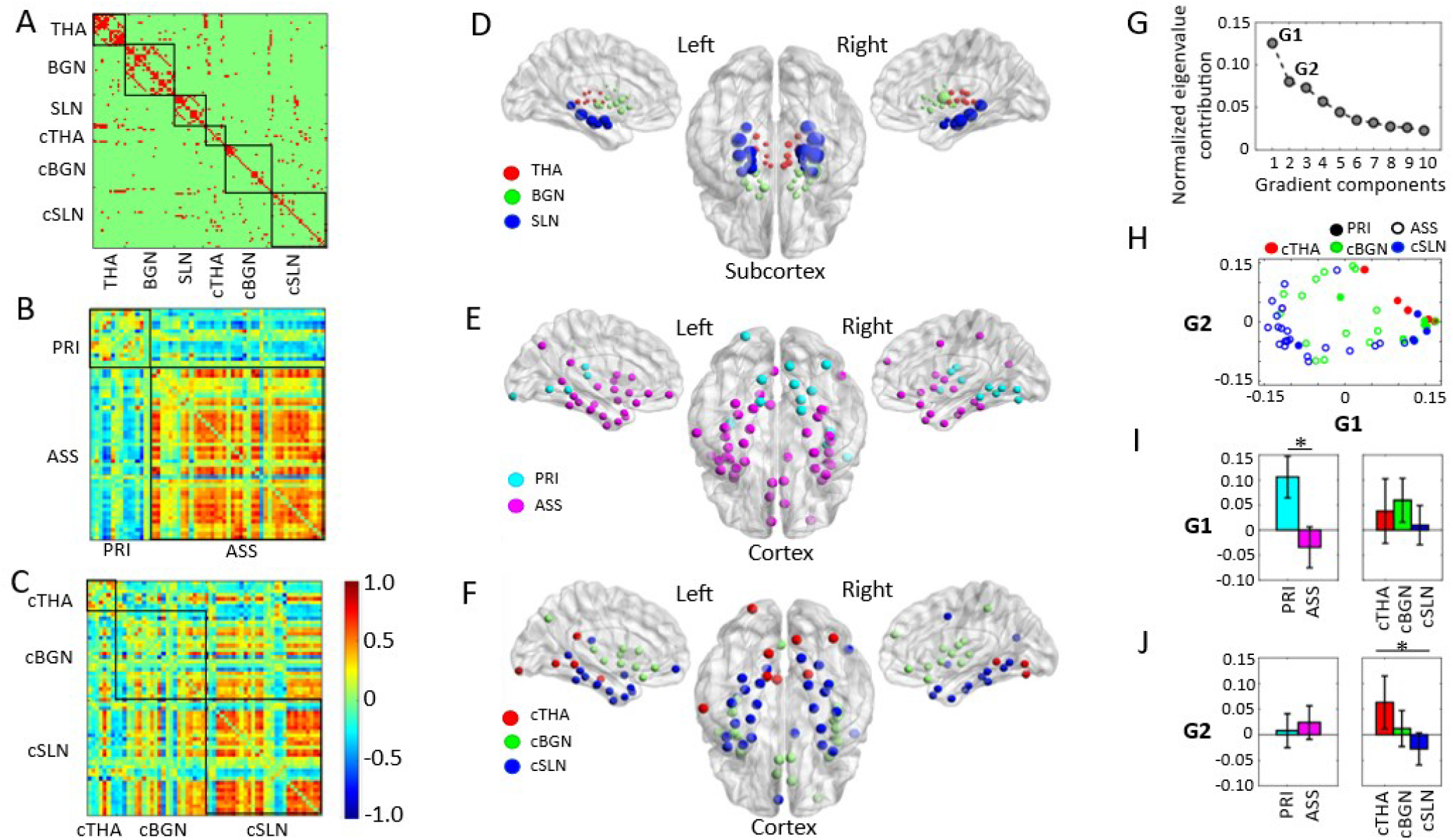
Functional convergence and microstructural organization of cortical regions converging on subcortical targets. (A) Group-level binary resting-state functional connectivity within subcortical target communities THA, BGN, and SLN and with their converging cortical regions (cTHA, cBGN, cSLN). (B–C) Convergence-restricted MPC matrices ordered by cortical type (PRI/ASS) and subcortical target, respectively; color bar indicates normalized MPC values. (D) Spatial distribution of subcortical target regions; node size represents the number of cortical regions (0 to 6) showing significant functional convergence with each subcortical region (created using BrainNet Viewer [12]). (E–F) Spatial distribution of convergence regions by cortical type and target, respectively. (G) Normalized eigenvalue contributions of microstructural gradients. (H) Distribution of cortical regions in G1–G2 space; filled/open circles represent PRI/ASS and colors represent cTHA/cBGN/cSLN. (I–J) Effects of *CortexType* and *Target* on G1 and G2, respectively. * indicates significant effects/comparisons (Bonferroni-corrected *p* < 0.05).

The corresponding convergence-restricted MPC matrices showed structured patterns of microstructural similarity when the regions were ordered according to cortical type (PRI/ASS; **Fig. 1B**) and according to their subcortical target (cTHA/cBGN/cSLN; **Fig. 1C**). The spatial distributions of these cortical regions according to cortical type and subcortical target are shown in **Fig. 1E** and **Fig. 1F**, respectively.

### 3.2. Microstructural gradient organization

Diffusion-map embedding of the convergence-restricted group-level MPC matrix yielded a series of low-dimensional microstructural gradients. The normalized eigenvalue contribution decreased across gradient components, with the largest contributions observed for G1 and G2 (**Fig. 1G**). These two gradients were therefore retained for subsequent analyses. The distribution of convergence-related cortical regions in G1–G2 space revealed complementary organization across the two dimensions, with PRI/ASS classification represented by marker fill and subcortical target represented by marker colour (**Fig. 1H**).

Each gradient was analysed separately using a two-way ANOVA with *Target* (cTHA, cBGN, cSLN) and *CortexType* (PRI, ASS) as factors, including the *Target* × *CortexType* interaction. For G1, there was a significant main effect of *CortexType*, with PRI regions showing higher G1 values than ASS regions (**Fig. 1I**; *F*(1,51) = 23.28, *p* < 0.001). The main effect of *Target* was not significant (*F*(2,51) = 1.45, *p* = 0.244), and the *Target* × *CortexType* interaction was also not significant (*F*(2,51) = 0.59, *p* = 0.56). For G2, the main effect of *Target* was significant (**Fig. 1J**; *F*(2,51) = 4.77, *p* = 0.01), whereas the main effect of *CortexType* (*F*(1,51) = 0.47, *p* = 0.50) as well as the interaction (*F*(2,51) = 0.22, *p* = 0.80) were not significant. Bonferroni-corrected pairwise comparisons indicated significant differences between cTHA and cSLN (*p* = 0.01). However, cBGN did not differ significantly from either cTHA (*p* = 0.32) or cSLN (*p* = 0.29), consistent with its intermediate proportion of primary and association cortical regions (32% PRI, 68% ASS), compared with cTHA (78% PRI, 22% ASS) and cSLN (18% PRI, 82% ASS).

For completeness, G3 and G4 were additionally examined using the same statistical analysis. For G3, only the main effect of *CortexType* was significant (F(1, 51) = 7.49, *p* = 0.01), whereas for G4, no significant effects of *Target, CortexType*, or their interaction were observed (all *p* > 0.22). Overall, the convergence-related cortical regions exhibited distinct dimensions of microstructural organization, with G1 and G3 primarily associated with cortical type and G2 showing target-related differentiation. No significant *Target* × *CortexType* interactions were observed across the analysed gradients.

### 3.3. Cytoarchitectural differences across cortical convergence regions

Cytoarchitectural composition showed that the *CortexType* × *CytoType* interaction was at the significance threshold, with primary and association convergence regions showing distinct overall cytoarchitectural profiles (**Fig. 2A**; *F*(5,255) = 2.24, *p* = 0.05). In contrast, the *Target* × *CytoType* interaction showed significance, indicating that the relative representation of cytoarchitectural types differed across cortical regions converging on THA, BGN, and SLN (**Fig. 2B**; *F*(5,255) = 2.39, *p* = 0.01). The three-way *CortexType* × *Target* × *CytoType* interaction was not significant (*F*(5,255) = 1.11, *p* = 0.35), indicating that the target-related cytoarchitectural differences were not significantly dependent on whether regions were classified as primary or association cortex. For visualization of target-related patterns within each cortical type, the profiles are shown in **Fig. 2C–D**. All other effects were nonsignificant (all *p* > 0.10).

**Fig. 2.**
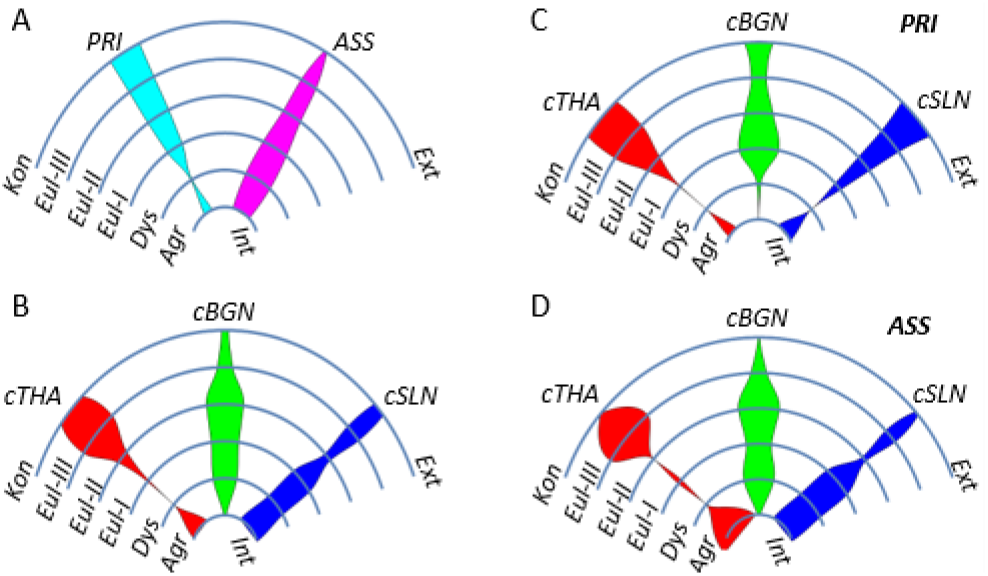
Cytoarchitectural composition of cortical convergence regions. Circular plots showing the mean percentage composition of six cytoarchitectural types [8]: koniocortical (Kon), eulaminate (Eul-III, Eul-II, Eul-I), dysgranular (Dys), and agranular (Agr), across cortical convergence regions. The percentage of each type is represented by the length of the respective curved line falling within the patch and not by the area between the curved lines covered by the patch [5]. Mean cytoarchitectural composition is shown for: (A) primary (PRI) and association (ASS) convergence regions, combined across targets; (B) cortical regions converging on THA, BGN, and SLN (cSUB), combined across cortical types; and (C–D) PRI and ASS convergence regions, respectively, separated by subcortical target. External (Ext) denotes the pial (outer) surface of the cortex, whereas internal (Int) denotes the white-matter boundary.

Bonferroni-corrected comparisons showed higher Kon in PRI than ASS regions (*p* < .001) and higher Dys in ASS than PRI regions (*p* = .04). Target-wise post-hoc comparisons further indicated higher Eul-III representation in THA-than BGN-converging regions (*p* = .01), higher Eul-II representation in BGN-than SLN-converging regions (*p* = .05), and higher Agr representation in SLN-than BGN-converging regions (although weaker at p = 0.08). Overall, the target-specific convergence patterns were characterized by distinct dominant cytoarchitectural types, with Eul-III, Eul-II, and Agranular predominating for THA, BGN, and SLN, respectively.

## 4. CONCLUSION

Our results demonstrate that cortical regions functionally converging on distinct subcortical communities exhibit systematic microstructural organization. Distinct gradient dimensions captured complementary aspects of this organization: G1 was primarily associated with primary-to-association cortical differentiation, whereas G2 showed differentiation related to subcortical target beyond cortical-type composition. Complementing these continuous microstructural gradients, cytoarchitectural analysis revealed target-related differences in the composition of cortical convergence regions, with distinct cytoarchitectural profiles predominating across THA, BGN, and SLN convergence. Together, these findings indicate that cortico-subcortical functional convergence is associated with multiple, complementary dimensions of cortical organization, encompassing both broad cortical hierarchy and target-related microstructural differentiation.

## 5. COMPLIANCE WITH ETHICAL STANDARDS

This study used de-identified human MRI data from the publicly available MICA-MICs dataset. The original study obtained approval from the relevant institutional ethics committee and written informed consent from all participants [4]. The present study involved secondary analysis of these publicly available data.

## 6. ACKNOWLEDGMENTS

No funding was received for conducting this study. The authors have no relevant financial or non-financial interests to disclose. We acknowledge the use of open AI tools for language refinement.

## 7. AUTHOR CONTRIBUTIONS

Conceptualization and Methodology, PV and SD; Data curation, SS; Formal analysis, PV and SS; Writing—original draft, PV; Writing—review and editing, all authors.

